# Multiple Exposures, Reinfection, and Risk of Progression to Active Tuberculosis

**DOI:** 10.1101/398271

**Authors:** Sarah F. Ackley, Robyn S. Lee, Lee Worden, Erin Zwick, Travis C. Porco, Marcel A. Behr, Caitlin S. Pepperell

## Abstract

A recent study reported on a tuberculosis outbreak in a largely Inuit village. Among recently infected individuals, exposure to additional active cases was associated with an increasing probability of developing active disease within a year. Using binomial risk models, we evaluated two potential mechanisms by which multiple infections during the first year following initial infection could account for increasing disease risk with increasing exposures. In the *reinfection model*, multiple exposures have an independent risk of becoming an infection, and infections contribute independently to active disease. In the *threshold model*, disease risk follows a sigmoidal function with small numbers of exposures conferring a low risk of active disease and large numbers of exposures conferring a high risk. To determine the dynamic impact of reinfection during the early phase of infection, we performed simulations from a modified Reed-Frost model of TB dynamics following spread from an initial number of cases. We parameterized this model with the maximum likelihood estimates from the reinfection and threshold models in addition to the observed distribution of exposures among recent infections. We find that both models can plausibly account for the observed increase in disease risk with increasing exposures, but the threshold model confers a better fit than a nested model without a threshold (p=0.04). Our simulations indicate that multiple exposures during this critical time period can lead to dramatic increases in outbreak size. In order to decrease TB burden in high-prevalence settings, it may be necessary to implement measures aimed at preventing repeated exposures, in addition to preventing primary infection.

## Introduction

While record low tuberculosis (TB) rates have been achieved in some countries, significant challenges in global TB control remain. In Canada, TB disease rates are considerably higher in the Inuit population than in the non-aboriginal Canadian-born population (1). A recent study reported on a TB outbreak in a largely Inuit village, in which 34 out of 149 individuals (23%) progressed to active disease within a year following recent infection (2). This risk is much higher that the oft cited 2–5% risk of progression following recent infection (3–5). The study indicated that, among recently infected individuals, exposure to additional active cases was associated with a significantly increased probability of developing active disease within a year. Previous work has discussed the plausibility (6) and potential impact of reinfection (defined in (7) as occurring 5 years or more after an initial infection) on TB dynamics (8). However, the dynamic impact of numerous, tightly spaced exposures and attendant infections has not been explored. Using binomial risk models (9) and data from (2), we evaluated two plausible mechanisms by which reinfection during this early, critical period can account for the increased disease risk: an independent effect of each infection on risk of progression (*reinfection model*) and a high risk of progression once the cumulative number of exposures exceeds a threshold value (*threshold model*). Contacts of TB cases producing aerosols with high burdens of *Mycobacterium tuberculosis* are more likely to become cases (10–12), suggesting an inoculum effect as a plausible biological mechanism for both these models. Both models are consistent with the observed data, though neither fully accounts for the observed high risk of active disease in this population following an initial exposure. We further find that multiple exposures during the critical, early time period following infection can dramatically increase expected outbreak size. The effects of delays in outbreak control may consequently be magnified by high rates of progression among multiply exposed individuals, with the potential to worsen outcomes for communities with limited access to health care resources.

## Methods

Based on previously published TB models (8), we developed binomial risk models for TB infection during the first year after infection and fit these models to data from a TB contact investigation in a largely Inuit (>90%) village of 933 people. Study design and data collection are described in detail elsewhere (2,13). Data consisted of the number of TB exposures for 149 recently-infected individuals, and whether these individuals developed active TB disease within the following year.

Model equations are given in the Supplemental Information. In the *reinfection model*, each exposure has an independent risk of becoming an infection, and each infection independently contributes to the risk of active disease. This model is an extension of models of reinfection occurring years after initial infection, now applied to the classic incubation period for primary TB disease (7,14,15). In the *threshold model*, a small number of exposures confers a low risk of active disease, while a large number confers a high risk of active disease and is modeled using a sigmoidal function. To statistically test for the presence of a threshold effect, we compare the threshold model to the *increasing-risk model*, a model nested within the threshold model with one fewer parameter. This model is constrained to have a fixed inflection point at zero exposures and the increase in risk of disease with increasing exposures is similar to that of the *reinfection model*. Using maximum likelihood estimation, we estimated key parameters governing TB progression for these three models. The observed Fisher information was used to determine confidence intervals for logit-(reinfection model) and log-transformed (threshold and increasing-risk models) parameters and the chi-squared likelihood ratio test was used to compare the threshold and increasing risk models in order to evaluate for the presence of a threshold effect. Chi-squared goodness-of-fit p-values were also calculated.

To determine the dynamic impact of reinfection within a year of initial infection, we performed simulations from modified Reed-Frost models (16) for two years following an initial influx of cases. We limited the simulations to two years, as it seemed likely that more aggressive control measures would be implemented for large outbreaks well within this timeframe. We parameterized these models with the maximum likelihood estimates from the reinfection and threshold models. We generated a number of exposures for each uninfected individual that approximated the observed distribution of exposures. Additional details and alternative parameterizations are given in the Supplemental Information. For the reinfection model, exposed individuals acquired a total number of infections *i* with probability ζ per exposure and progressed to active disease within a year with probability 1-(1-p)^*l*^, where p is the risk of active disease for a single infection. For the threshold model, risk of disease was calculated from the maximum likelihood parameter estimates and equation given in the Supplemental Information. For both models, exposures and infections were generated for a second year using the number of cases produced in the first year. It was assumed all infectious individuals received treatment and were no longer infectious for a second year. The total cases over the two years, not including the initial cases, were then summed to obtain the outbreak size. We compared the threshold and reinfection models to a *constant risk model* parameterized from the reinfection model. In this model, risk of infection was a constant 12% for infected individuals regardless of the number of exposures, corresponding to the risk of disease for singly exposed infected individuals in the reinfection model. We investigated scenarios ranging from a single initial case to 50 initial cases.

## Results

Selected parameter estimates and confidence intervals for each of the three models are shown in Table 1. The remainder of the parameter estimates are in the Supplemental Information. Figure 1 shows the observed risk of infection as a function of number of exposures with binomial confidence intervals, as well as each of the model fits. Based on their goodness-of-fit p-values, the reinfection, increasing-risk, and threshold models are consistent with the observed data. The threshold model confers an improved fit over the increasing-risk model (p=0.04), providing evidence of a threshold effect. All models estimate a probability of progression to disease within one year for a single infection of over 10%, and significantly greater than 5%. The inflection point in our threshold model is at 17.6 exposures, indicating that the probability of active disease was larger for individuals with 18 or more exposures than for individuals with 17 or fewer exposures. Simulation from our modified Reed-Frost models indicates that allowing for reinfection or threshold effects produces larger outbreak sizes on average than a constant-risk model. The reinfection model increasingly diverges from the constant-risk model with increasing numbers of initial cases. The threshold and reinfection models produce outbreaks of similar size, which are larger than those produced in a constant-risk model. These results are summarized in Figure 2.

**Table 1:** Parameter estimates for the reinfection and threshold models. Confidence intervals are given in parentheses. A complete list of parameters for the threshold model is given in the Supplemental Information.

| Quantity | Value |  |  |
| --- | --- | --- | --- |
|  | Reinfection Model | Threshold Model | Increasing-Risk Model |
| *Negative log-likelihood* | 73.2 | 70.8 | 72.8 |
| *Probability of progression to disease (p)* | 0.12<br>(0.060, 0.21) | — | — |
| *Probability of infection given exposure ($\zeta$)* | 0.18<br>(0.055, 0.45) | — | — |
| *Probability of progression to disease given one exposure* | — | 0.15<br>(0.093, 0.24) | 0.12<br>(0.064, 0.22) |
| *Threshold location* | — | 17.7<br>(13.9, 22.5) | Constrained at 0 |
| *Increase in probability of progression to disease associated with many exposures* | — | 0.45<br>(0.27, 0.75) | — |
| *Goodness-of-fit p-value* | 0.97 | 0.99 | 0.97 |
| *P-value for significantly improved fit of threshold model* | — | 0.04<br>($\chi^2$ likelihood ratio test) | |

**Figure 1:**
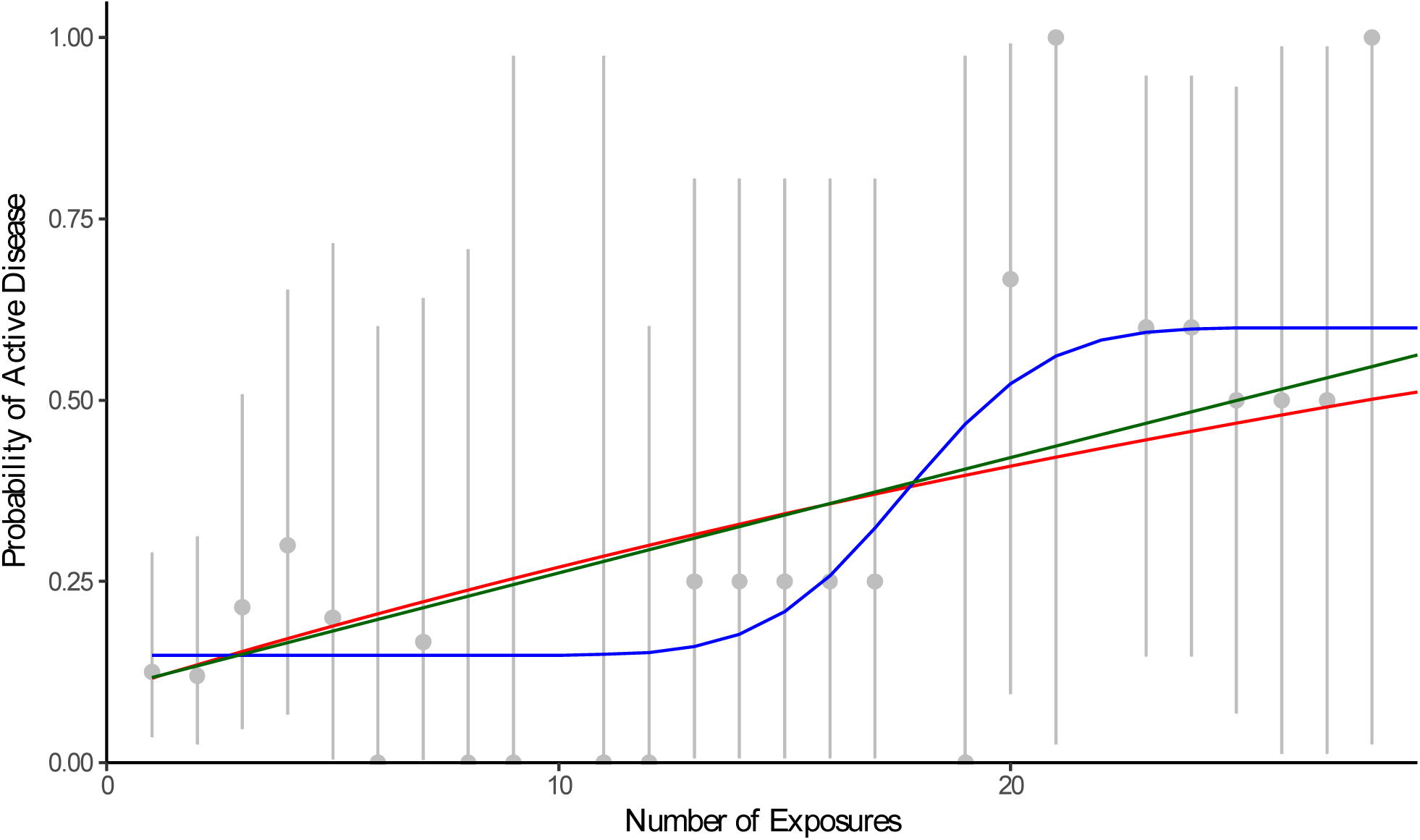
Data and model fit. The reinfection, threshold, and increasing-risk models are shown in red, blue, and green, respectively. Observed data are shown in grey with exact binomial confidence intervals.

**Figure 2:**
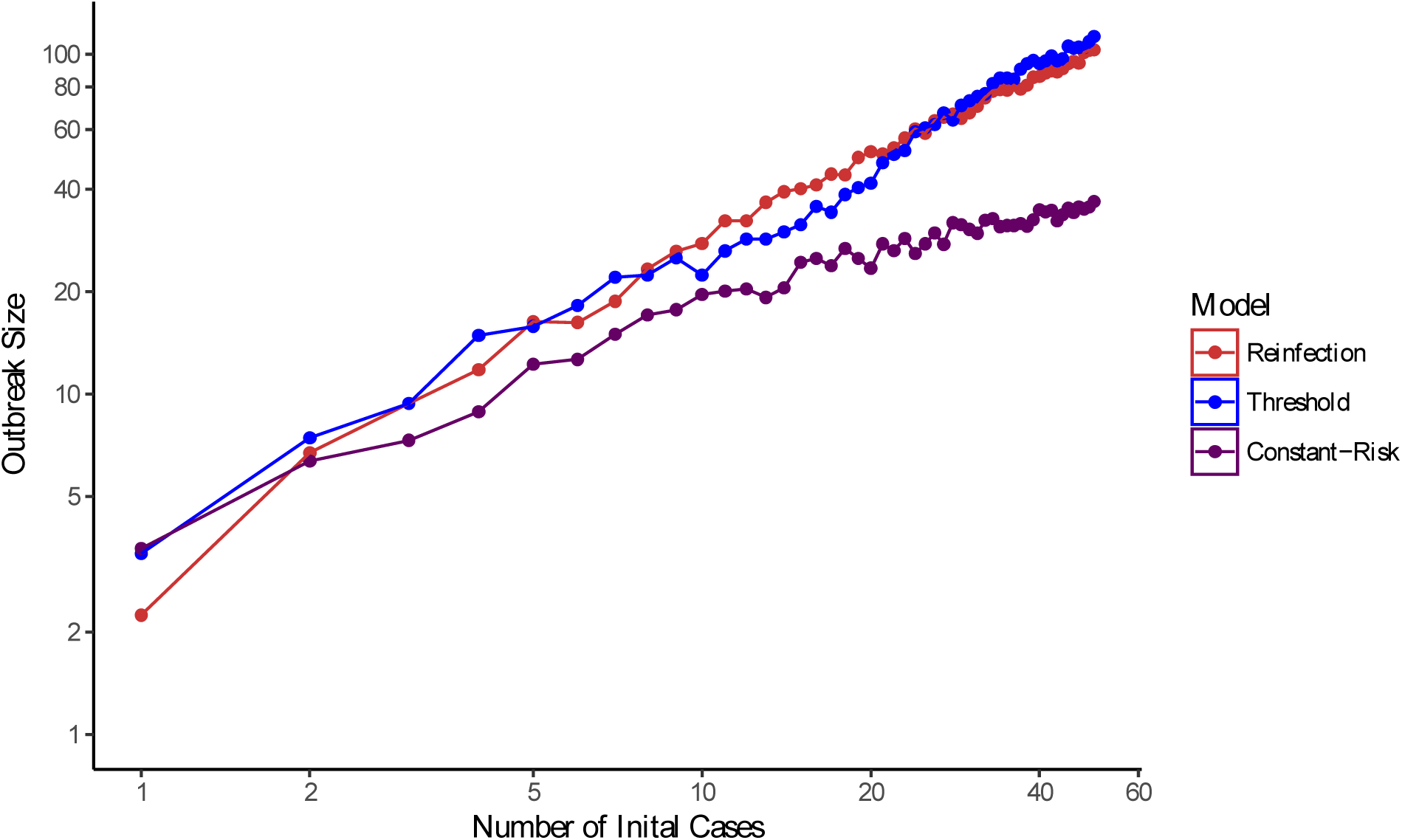
Expected outbreak size as a function of number of initial cases for the reinfection model, threshold model, and constant-risk model. Shaded ribbons depict the 95% confidence interval for the expected outbreak size. Twenty-five simulations were performed for each number of index cases, 1 to 50. Note that the individual risk of active disease is on average larger for the reinfection and threshold models than it is for the constant-risk model; for the constant-risk model, it is assumed that risk of disease from singly exposed infected individuals can be extrapolated to individuals with more exposures, a common practice in TB modeling.

## Discussion

The increase in disease risk with increasing exposures found in (2) could be explained by an independent effect of each infection on risk of progression (*reinfection model*), or by a high risk of progression once the cumulative number of exposures exceeds a threshold value (*threshold model*). Previous work has shown that the effect of increasing disease risk with increasing exposures persists after adjustment for a number of covariates (2). Here we found that the threshold model conferred an improved fit relative to the increasing-risk model, providing evidence for a threshold effect. However, more research is needed to determine whether this finding is replicable across human populations, and whether there is evidence of a threshold effect in other settings.

While reinfection or threshold effects can account for approximately half of the observed 20% risk of disease following recent infection in this population, it cannot account for all of it: all of our models estimated a probability of progression to active disease within a year given a single infection of over 10%, which is much higher than has been observed in other populations (17) and significantly greater than 5% (3–5). Various factors could contribute to this discrepancy: First, number of contacts may have been underreported for individuals with small numbers of exposures. Second, the nature of contacts between infectious and susceptible individuals in this setting may have resulted in a high rate of progression to disease among contacts. Intensity of exposure to an infectious case has been shown to affect the risk of TB infection and disease (18,19). Third, prior exposure to *Mycobacterium tuberculosis* or prior TB disease may have modified disease risk for individuals with few exposures. While prior infections are generally thought to protect against new infections (17,20,21), a South African study found prior disease to be associated with a fourfold *increased* risk of developing active TB following reinfection (22). Finally, closely spaced infections may act to modulate immune responses in a way that increases the risk of disease and transmissibility. Recent research supports the idea that multiple closely spaced exposures have a causal effect on disease risk—e.g. a recent study in the rabbit model of TB found that a bacterial inoculum divided over multiple exposures produced more extensive cavitary pulmonary disease than when the same inoculum was given in a single exposure (23). Additional information on the timing of infections and prior disease for individuals in this community and the timescales over which protective immunity develops for both individuals (with and without prior TB disease) would be required to determine how interactions among TB infections could have contributed to the observed high risk of disease.

Our simulation results indicate that reinfection and/or threshold effects can have a dramatic effect on outbreak sizes, even on short time scales. In situations where we expect a lag in intervention due to lack of resources or access to care, the probability of uninfected individuals coming into contact with multiple TB cases becomes increasingly likely. This suggests that delays in the implementation of outbreak control measures could contribute to health disparities; not only would more transmission occur during such delays, but increased exposure would also lead to significantly more active disease among those already exposed. These effects would propagate, ultimately leading to significantly higher disease burden than would be expected without such effects.

Failure to account for the effect of multiple exposures in models of TB might lead to spurious results. For example, the risk of active disease following infection may not be equivalent between low and high prevalence settings; this could lead to dramatic underestimation of disease burden in high prevalence settings, where multiple exposures are likely common. Conversely, fitting TB models to data from high prevalence settings might lead researchers to incorrectly conclude that the observed high risk of disease in a population is due to shared risk behaviors or high genetic susceptibility to disease, and thus underestimate the importance of early intervention in disease prevention at the population level. In terms of the practical implementation, we note that both the reinfection and threshold models predict similar outbreak sizes over two years over a wide range of initial cases, indicating that multiple parameterizations of the increasing risk of active disease with increasing exposures may be able to produce comparable outbreak sizes.

A limitation of our results is that numbers of exposures are based on self-reported contacts, which were then used to obtain the number of TB exposures for each infected individual. Underreporting of contacts could have contributed to the observed high disease rates and would lead to over estimates of the effect of each additional exposure. In addition, while we examined the effect of multiple closely spaced exposures on disease risk and consequent total outbreak sizes, multiple exposures may contribute to large outbreak sizes by other mechanisms as well, which could worsen the dynamic impacts of repeated exposures. Studies in humans have long indicated variability in infectiousness, with recent work suggesting that cough aerosols produced by a subset of TB patients enhance transmission (10,11). As noted above, experimental data suggest that individuals with multiple closely spaced infections could plausibly be at risk of developing more extensive/infectious forms of TB and thereby delivering large bacterial inocula associated with a high risk of disease progression in their contacts. If multiply exposed individuals produce higher levels of aerosols, this could further amplify the effects of closely spaced infections and attendant impacts of delays in treatment and case finding. Further work would be required to determine the effect of multiple closely spaced exposures on disease risk.

Theoretical work has indicated that exogeneous reinfection occurring years after initial infection can lead to unpredictable TB dynamics (e.g. multiple equilibria), which in turn can decrease the effectiveness of public health interventions (8). We find here that reinfection within a year of initial infection can lead to appreciably larger outbreak sizes. TB models for high prevalence settings may need to account for this in order to obtain unbiased estimates of the effectiveness of interventions. It may also be essential to implement measures aimed at preventing repeated exposures, in addition to preventing primary infection, in order to decrease TB burden in high-prevalence populations. Early intervention and access to health care are essential to addressing disparities in TB burden.

## Ethics

This study was a secondary analysis of previously collected data. Data for this study was completely deidentified. The only variables used were number of exposures and case status.

## Data accessibility

Data and code are available as part of the electronic supplementary information.

## Authors’ contributions

CSP developed the research question and RSL and MAB contributed relevant data. SFA, TCP, an CSP developed an analysis plan. SFA, LW, and EZ performed the analysis. SFA, RSL, MAB, and CSP interpreted the results and wrote the manuscript. All authors give final approval for publication.

## References

1. Public Health Agency of Canada. The Chief Public Health Officer’s Report on the State of Public Health in Canada 2013 – Tuberculosis – past and present [Internet]. AEM. 2013 [cited 2017 Nov 19]. Available from: https://www.canada.ca/en/public-health/

2. Lee RS, Proulx J-F, Menzies D, Behr MA. Progression to tuberculosis disease increases with multiple exposures. Eur Respir J. 2016 Oct 6;ERJ-00893-2016.

3. Mack U, Migliori GB, Sester M, Rieder HL, Ehlers S, Goletti D, et al. LTBI: Latent tuberculosis infection or lasting immune responses to M. tuberculosis? A TBNET consensus statement. Eur Respir J. 2009 May 1;33(5):956–73.

4. Sloot R, Schim van der Loeff MF, Kouw PM, Borgdorff MW. Risk of Tuberculosis after Recent Exposure. A 10-Year Follow-up Study of Contacts in Amsterdam. Am J Respir Crit Care Med. 2014 Sep 29;190(9):1044–52.

5. Downes J. A Study of the Risk of Attack among Contacts in Tuberculous Families in a Rural Area. Am J Hyg. 1935;22:731–42.

6. Cardona P-J. A Dynamic Reinfection Hypothesis of Latent Tuberculosis Infection. Infection. 2009 Apr 1;37(2):80.

7. Vynnycky E, Fine PE. Interpreting the decline in tuberculosis: the role of secular trends in effective contact. Int J Epidemiol. 1999 Apr;28(2):327–34.

8. Feng Z, Castillo-Chavez C, Capurro AF. A model for tuberculosis with exogenous reinfection. Theor Popul Biol. 2000 May;57(3):235–47.

9. Porco TC, Martin JN, Page-Shafer KA, Cheng A, Charlebois E, Grant RM, et al. Decline in HIV infectivity following the introduction of highly active antiretroviral therapy. AIDS Lond Engl. 2004 Jan 2;18(1):81–8.

10. Jones-López EC, Namugga O, Mumbowa F, Ssebidandi M, Mbabazi O, Moine S, et al. Cough aerosols of Mycobacterium tuberculosis predict new infection: a household contact study. Am J Respir Crit Care Med. 2013 May 1;187(9):1007–15.

11. Jones-López EC, Acuña-Villaorduña C, Ssebidandi M, Gaeddert M, Kubiak RW, Ayakaka I, et al. Cough Aerosols of Mycobacterium tuberculosis in the Prediction of Incident Tuberculosis Disease in Household Contacts. Clin Infect Dis Off Publ Infect Dis Soc Am. 2016 Jul 1;63(1):10–20.

12. Jones-López EC, Acuña-Villaorduña C, Fregona G, Marques-Rodrigues P, White LF, Hadad DJ, et al. Incident Mycobacterium tuberculosis infection in household contacts of infectious tuberculosis patients in Brazil. BMC Infect Dis. 2017 Aug 18;17.

13. Lee RS, Radomski N, Proulx J-F, Manry J, McIntosh F, Desjardins F, et al. Reemergence and Amplification of Tuberculosis in the Canadian Arctic. J Infect Dis. 2015 Jun 15;211(12):1905–14.

14. Ackley SF, Liu F, Porco TC, Pepperell CS. Modeling historical tuberculosis epidemics among Canadian First Nations: effects of malnutrition and genetic variation. PeerJ. 2015 Sep 24;3:e1237.

15. Castillo-Chavez C, Song B. Dynamical models of tuberculosis and their applications. Math Biosci Eng MBE. 2004 Sep;1(2):361–404.

16. Abbey H. An Examination of the Reed-Frost Theory of Epidemics. Hum Biol. 1952 Sep 1;24(3).

17. Vynnycky E, Fine PE. The natural history of tuberculosis: the implications of age-dependent risks of disease and the role of reinfection. Epidemiol Infect. 1997 Oct;119(2):183–201.

18. Acuña-Villaorduña C, Jones-López EC, Fregona G, Marques-Rodrigues P, Gaeddert M, Geadas C, et al. Intensity of exposure to pulmonary tuberculosis determines risk of tuberculosis infection and disease. Eur Respir J. 2018 Jan 1;51(1):1701578.

19. Lutong L, Bei Z. Association of prevalence of tuberculin reactions with closeness of contact among household contacts of new smear-positive pulmonary tuberculosis patients [Notes from the Field]. Int Union Tuberc Lung Dis. 2000 Mar;4(3):275–7.

20. Ferguson RG. Tuberculosis Among the Indians of the Great Canadian Plains. London: Adlard and Son; 1928.

21. Brooks-Pollock E, Becerra MC, Goldstein E, Cohen T, Murray MB. Epidemiologic Inference From the Distribution of Tuberculosis Cases in Households in Lima, Peru. J Infect Dis. 2011 Jun 1;203(11):1582–9.

22. Verver S, Warren RM, Beyers N, Richardson M, van der Spuy GD, Borgdorff MW, et al. Rate of reinfection tuberculosis after successful treatment is higher than rate of new tuberculosis. Am J Respir Crit Care Med. 2005 Jun 15;171(12):1430–5.

23. Urbanowski ME, Ihms EA, Bigelow K, Kübler A, Elkington PT, Bishai WR. Repetitive Aerosol Exposure Promotes Cavitary Tuberculosis and Enables Screening for Targeted Inhibitors of Extensive Lung Destruction. J Infect Dis.

